# Coupling hydrodynamic drifting simulations and seasonal demographics to forecast the occurrence of jellyfish blooms

**DOI:** 10.1101/2024.03.11.584451

**Authors:** James Cant, Owen R. Jones, Ingrid Ellingsen, Jack H. Laverick, Sanna Majaneva, Jan Dierking, Nicole Aberle, Jamileh Javidpour

## Abstract

1. Although jellyfish are an important component of coastal marine communities, their public perception is often tainted by their proclivity for aggregating in vast numbers, known as jellyfish blooms. Jellyfish blooms occur worldwide and are associated with major economic ramifications, particularly throughout the fisheries, aquaculture, and tourism sectors.
2. Forecasting jellyfish blooms is critical in order to manage and mitigate their ecological and economic impacts, but the complex life cycles and cryptic life stages exhibited by most jellyfish species largely precludes accurate predictions of their temporal and spatial occurrence.
3. Here we introduce a predictive framework, combining state-of-the-art hydrodynamic simulations and periodic population modelling approaches, to forecast spatial and temporal patterns in the formation of jellyfish blooms. While this framework is sufficiently flexible for accommodating various bloom-forming jellyfish species and impacted coastal regions worldwide, we focus on moon jellyfish (*Aurelia aurita*) populations within the Baltic Sea as an illustrative example.
4. We emphasise how, through the iterative process of forecasting and model validation, this framework can provide valuable insights for resolving key gaps in our understanding of the drivers of bloom events. Indeed, our framework offers an approach for identifying the, to date, unknown locations of polyp-beds; a crucial parameter in enhancing our capacity to accurately predict the occurrence of jellyfish blooms. Accordingly, our framework, represents a key decision-support tool for mitigating the socioeconomic impacts of bloom formation.

## 1. Introduction

The intensifying need for adaptive, rather than reactive, strategies to prevent biodiversity loss challenges ecologists to forecast the outcome of complex interactions between natural populations and their environments (Mouquet et al., 2015; Petchey et al., 2015). Yet, with ongoing global change rarely affording enough time to acquire sufficient data for making these predictions, there is a reluctance to use forecasting tools for fear of inaccuracy (Houlahan et al., 2017). Ecological forecasting is a balance of science and art, reliant on rigorous mathematical and statistical methods, but underpinned by ‘artistic’ choices around the modelling approach to use, the elements to include (and ignore in the interest of parsimony), and how to address underlying model assumptions (Tetlock & Gardner, 2015). However, while we must acknowledge the impacts of uncertainty on our capacity to predict future events, it should not inhibit ecological forecasting (Clark et al., 2001). Making predictions, and then validating them, offers the opportunity for identifying the ecological characteristics we do not yet understand, thus providing insight into where additional research is needed (Carpenter, 2002; Houlahan et al., 2017; Merow et al., 2014). Indeed, using iterative forecasting tools to first highlight and then resolve gaps in our ecological understanding is just as valuable as their use in anticipating ecological events (Dietze & Lynch, 2019).

As ecology seeks to become a more predictive science (Dietze et al., 2018; Johnson et al., 2021), understanding the spatial movements of natural populations, and their abiotic cues, is becoming a key management and conservation tool (McLeod & Leroux, 2021). The localized and migratory movements of organisms significantly impact their vulnerability to temporal and spatial shifts in climatic conditions and recurrent disturbances (Robinson et al., 2009; Schloss et al., 2012). In light of this, current discourse should pivot towards developing climate-adaptive management strategies, enabling the forecasting of ecological population changes. Accounting for the spatial dynamics of populations can also help to resolve the deficiencies associated with short ecological time series (Johnson et al., 2021). Moreover, an explicit appreciation for how organisms utilise their local landscapes can help us to manage the socioeconomic and ethical impacts of human-wildlife conflicts (Kremen & Merenlender, 2018; Reynolds et al., 2017). However, a major challenge associated with predicting the spatial movements of natural populations is that, for many species, key aspects of the complex interaction between the physiological tolerances and dispersal pathways of individual organisms remain cryptic.

Jellyfish are key components of marine biodiversity and fulfil a diverse range of ecological roles, contributing to intricate food web networks, nutrient cycling, and carbon sequestration (Chi et al., 2018; Choy et al., 2017; Hays et al., 2018; Wright et al., 2021). Yet, beyond their many ecological benefits, the sudden appearance of vast numbers of individuals in so-called jellyfish blooms can also be problematic (Brodeur et al., 1999; Lotan et al., 1994; Uye, 2008). These bloom events, which occur following increased reproduction and growth (true blooms) or the localised accumulation of multiple populations (apparent blooms; Graham et al., 2001; Hamner & Dawson, 2009), influence ecosystem functioning, and can be detrimental to fishing, aquaculture, and tourism activities (Gershwin et al., 2013; Graham et al., 2014). Moreover, while it is debated whether the causal mechanisms are increasing global jellyfish abundances or increased human exploitation of coastal environments, there is a growing perception that the recurrence frequency of jellyfish bloom events is increasing (Condon et al., 2013). Regardless of this apparent increase, however, the dynamics of jellyfish populations and thus the causal mechanisms of bloom formation remain poorly understood (Fernández-Alías et al., 2024; Goldstein & Steiner, 2020; Kennerley et al., 2021), leaving us unable to mitigate their broad ecological and economic impacts (Condon et al., 2013).

Our limited knowledge of jellyfish population dynamics arises due to their complex metagenic life cycles which consist of pelagic and benthic stages separated in both time and space (Lucas, 2001). These complex life cycles are further complicated by high stage-specific plasticity in response to changing environmental conditions (Goldstein & Steiner, 2020; Hirst & Lucas, 1998; Kennerley et al., 2021). Here, we introduce a predictive framework, combining hydrodynamic simulations and seasonal demographic patterns, that aims to enhance our understanding of the mechanisms governing the timing and geographical location of jellyfish blooms. As a proof of concept, we present this framework focusing on the dynamics of moon jelly (*Aurelia aurita* [Linnaeus, 1758], Scyphozoa) populations within the Baltic Sea. However, we discuss how the tool is sufficiently flexible for accommodating assessments of other species and settings as needed. Using an approach similar to demographic distribution modelling (*sensu* Merow et al., 2014), our framework combines hydrodynamic drifting simulations produced by a biogeochemical-ocean circulation model (SINMOD; Slagstad et al., 2011; Slagstad & McClimans, 2005; Wassmann et al., 2006) with periodic matrix population models (Bacaër, 2009; Caswell, 2001). We also illustrate how this framework presents a valuable tool for forecasting spatial and temporal patterns in jellyfish bloom occurrence and, crucially, for identifying key gaps in our understanding of the drivers of bloom events.

## 2. Materials & Methods

### 2.1. Species and study area

*Aurelia aurita* is a cosmopolitan jellyfish species and comprises one of the most abundant bloom-forming species within the Baltic region (Stoltenberg et al., 2021). Throughout their life cycle, *A. aurita* individuals pass through a series of discrete stages, each with differing vital rate profiles (Goldstein & Steiner, 2020; Fig. 1). Starting as sexually produced planula larvae, *A. aurita* individuals settle on suitable substrate to form polyps known as scyphistoma (Lucas, 2001). Although the polyp phase of *A. aurita* is highly cryptic, populations can achieve considerable densities as individuals propagate through asexual fission (Goldstein & Steiner, 2020; Lucas et al., 2012). Eventually, individual polyps undergo a form of transverse fission called strobilation, during which they release free-swimming ephyra (Lucas et al., 2012). These ephyra then develop into sexually mature adult medusae, the characteristic nomadic phase of jellyfish life cycles and, ultimately, the key bloom-forming stage (Lucas & Dawson, 2014).

**Figure 1.**
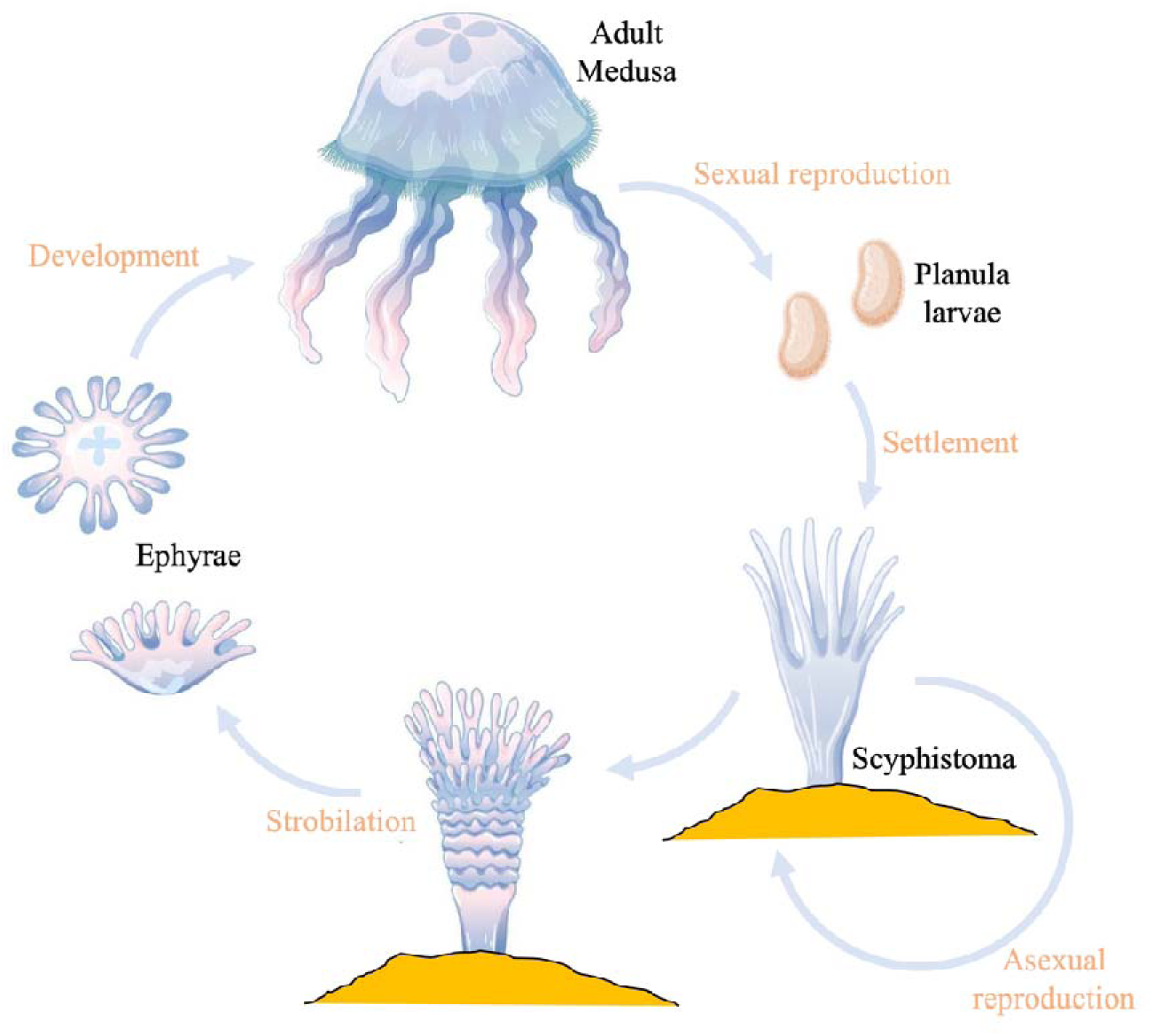
Schematic diagram reflecting the complex metagenic life cycle exhibited by *Aurelia aurita*, during which individuals transition between benthic asexually reproducing polyps, or scyphistoma, and sexually reproducing pelagic medusae.

Our study area, the Baltic Sea, is a relatively shallow, brackish body of water characterised by strong temperature and salinity gradients arising from the mixing of saltwater from the North Sea with riverine outflows from across Scandinavia (Ojaveer et al., 2010). Due to its young geological age and salinity constraints, the Baltic Sea is relatively species-poor, yet this region is an area of high research interest due to its exposure to oceanic warming, over-fishing, acidification, nutrient contamination, and deoxygenation (Reusch et al., 2018 and references therein).

### 2.2. Forecasting Medusae Dynamics

Our predictive framework consists of three analytical functions *DMat*, *JellySim*, and *SimPlot*, developed using *R* (R Core Team, 2019; all function scripts are available at https://github.com/CantJ/Bloom-forecasting-tools). Forecasting the appearance of jellyfish blooms requires understanding the development of adult medusae. Medusae development is contingent on the environmental conditions individuals experience as they drift through the water column, which depend on when and where the individuals are released from their polyp beds. Accordingly, our analytical functions comprise two key elements: (1) a demographic component describing how abiotic conditions influence the survival, development, and reproduction of individuals (*DMat*) and (2) a hydrodynamic drifting component that details the spatial movements expected of individuals and the environmental conditions they experience as a result (*JellySim*; Fig. 2). Meanwhile, *SimPlot* handles the plotting capabilities of our framework, returning the combined outputs of *DMat* and *JellySim* as a series of heat maps allowing users to visualise where, in both space and time, aggregations of medusae are most likely to occur given the abiotic and demographic parameters provided.

**Figure 2.**
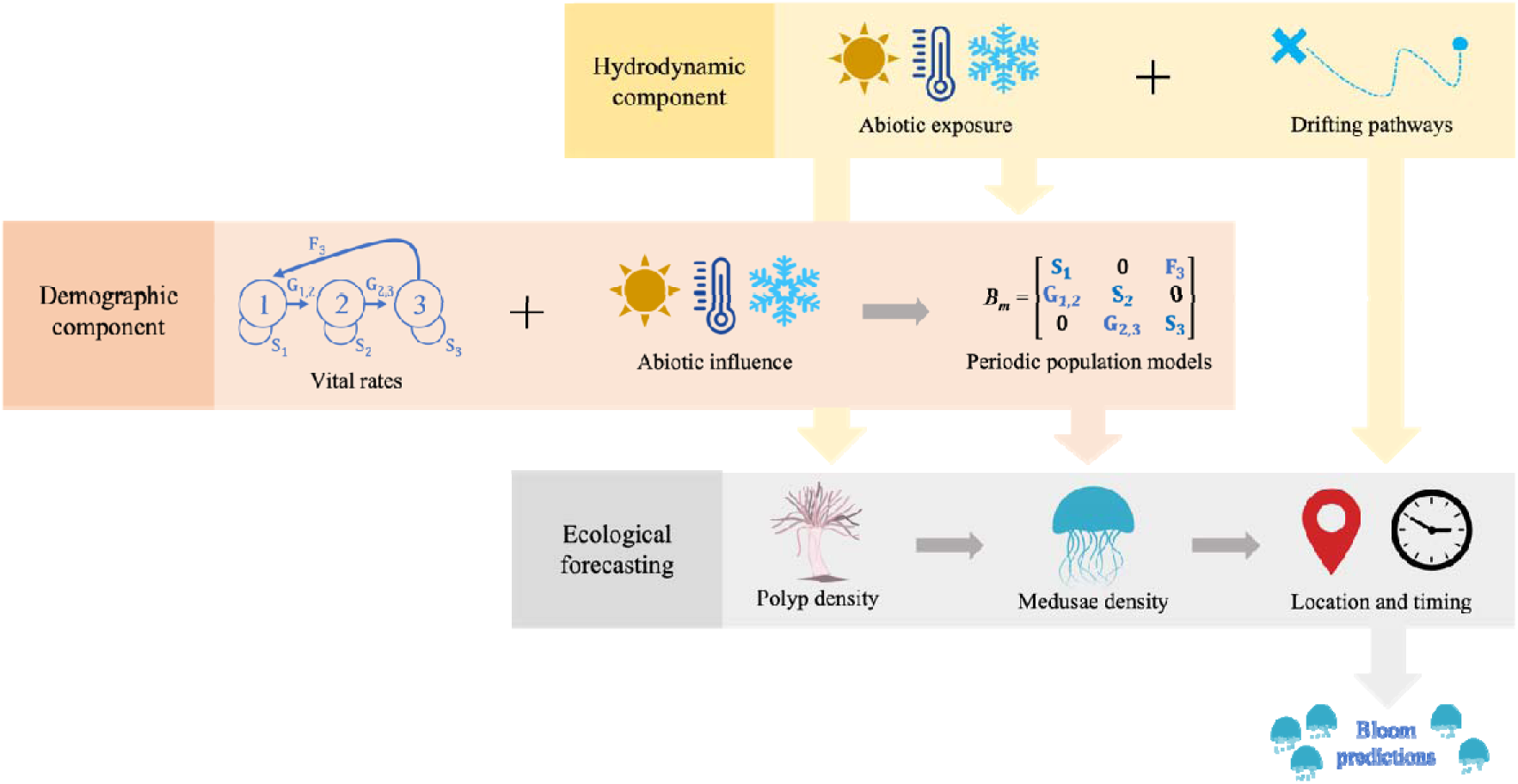
Our predictive framework combines hydrodynamic drifting simulations and seasonal demographic modelling to forecast spatial and temporal patterns in the occurrence of jellyfish blooms. The demographic component of this framework comprises the vital rate profiles of survival (S), growth (G) and reproduction (F) across the life cycle of *A. aurita*, and how these patterns vary in response to changing temperature and salinity regimes. By coupling these demographic profiles with particle track pathways obtained from hydrodynamic drifting simulations, and the environmental conditions experienced along these pathways, this framework is then able to forecast spatial and temporal patterns in the appearance and aggregation of *A. aurita* individuals within the Baltic Sea, and subsequently the likelihood of bloom events.

#### 2.2.1. Demographic model construction and parameterisation

The demographic component of our framework is handled by the *DMat* function which, when provided with details outlining local abiotic regimes, estimates an initial population of sessile polyps (polyp density, polyp cm^-2^) for the selected focal location(s), before quantifying the asexual dynamics of this polyp population, their release of ephyrae, and the subsequent development and survival of medusae. Here, we outline the structure of the demographic models contained within the *DMat* function and the approach used to parameterise these models required for our example *A. aurita* population in the Baltic Sea. However, we emphasise that one of the advantages of our framework is that, by including all of the demographic elements within a single function, it is possible for users to replace this function with alternative modelling approaches more appropriate for their focal population(s) and/or data availability. Provided these alternative functions are able to receive and provide inputs and outputs compatible with the *JellySim* function (see *2.2.2. Implementing forecasting*), then our framework will still be able to simulate the temporal and spatial movements of these populations.

To become medusae, jellyfish must successfully transition through a series of discrete life-stages (*i.e*., planula, scyphistoma, and ephyra; Fig. 1); a process mediated by the vital rates of survival, growth, and reproduction, and how these vital rates are influenced by environmental conditions (Goldstein & Steiner, 2020). Stage-structured matrix population models (MPMs; *sensu* Lefkovitch, 1965) summarise stage-specific survival, development, and reproduction, into a population projection matrix (***A***). Across this projection matrix, each element (a_ij_) describes the probability of individuals transitioning from stage *j* into stage *i* across the discrete time period *t* to *t+1*, or the per-capita reproductive contributions of state *j* individuals into state class *i* during that same interval (Caswell, 2001; Groenendael et al., 1988). This matrix can then be analysed to explore how population size (*N*, number of individuals), and structure (*n,* the proportion of individuals in each life-stage) are expected to change over time:

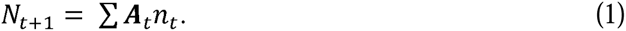

Accordingly, our *DMat* function is based on this matrix modelling approach to describe the transition of *A. aurita* individuals from larvae to medusae, through the scyphistoma and ephyra life-stages.

In *A. aurita* populations, once ephyrae metamorphose into medusae they are not immediately reproductively viable. Instead, juvenile medusae must grow beyond a certain size before they become sexually mature, a threshold which is heavily influenced by local environmental conditions (Lucas & Lawes, 1998; Lucas & Williams, 1994). Consequently, once *A. aurita* individuals become medusae, their dynamics and, ultimately, their ability to contribute to larval production, are best described on a continuous size scale. Therefore, following the transition of individuals into the medusa life-stage, our approach adopts a continuous integral projection model (IPM) framework. Fundamentally, IPMs are analogous to MPMs but, rather than being structured by discrete state variables, they allow populations to be structured according to a continuous state variable, such as size (Easterling et al., 2000). Hence, in our model, we opted to represent medusae dynamics using an IPM format structured by medusae size (bell diameter, mm).

Finally, to accommodate the highly seasonal nature of the *A. aurita* life-cycle, and to allow us to explore the within-year timing of bloom formation, *DMat* relies on a periodic population modelling approach (Bacaër, 2009; Caswell, 2001). Most matrix models use a projection interval (*t* to *t+1*) of 1 year (Caswell, 2001) corresponding with the commonplace yearly cycle of productivity and limited resource availability (Varpe, 2017), but this approach cannot capture within-year dynamics. In periodic population models, population matrix ***A***, and thus the time-frame it represents, is broken-down into *m* periodic matrices (***B***; Caswell, 2001), allowing users to better capture within-year episodic shifts across the vital rates of survival, development, and reproduction:

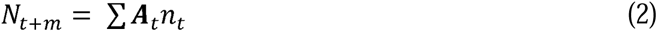

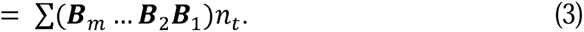

We used this approach to define a series of monthly periodic matrices, allowing us to explore fine-scale temporal patterns in the dynamics of *A. aurita* populations.

We estimated the state-specific transition rates for our demographic models using data available within the GoJelly databank (Volpe et al., 2023). Briefly, this database consists of records compiled from independent experimental and field observations of demographic patterns across various jellyfish species and life-stages (e.g., larval settlement probability and the number of ephyrae produced per polyp). Focusing only on entries for *Aurelia* spp., we extracted data subsets corresponding to the differing dynamics of planula larvae, scyphistomae, ephyrae, and medusae. With these data subsets, we used generalised linear mixed models (GLMMs) to quantify the characteristics underlying the stage-specific patterns of survival, growth, and reproduction, necessary for quantifying the complete life cycle of *A. aurita* (see Supplementary Material).

Firstly, we determined the vital rates associated with the transition (*i.e*., development and clonality), and persistence (*i.e.*, stasis) of *A. aurita* individuals across the discrete planula, scyphistoma, and ephyra phases of the life cycle (Table S1). We quantified each characteristic as the intercept of a fixed-slope GLMM, with temperature (°C) and/or salinity (ppt) included as fixed-effects (we note that for planula stasis and survival, and polyp asexual propagation, there were no data in the databank linking demographic and abiotic properties). Across all of our GLMMs, we modelled probabilistic characteristics (e.g., larval settlement probability, survival [all phases], strobilation, and ephyra metamorphosis) using logistic regression, while we modelled polyp density, and the mean number of ephyrae produced per polyp, using a gamma model. Using a gamma distribution ensured a continuous but non-negative and non-zero scale for both polyp density and mean ephyra production per polyp individual. Meanwhile, we modelled the number of individuals produced during asexual polyp propagation using a zero-adjusted gamma distribution (ZAGA). Adopting a ZAGA distribution enabled us to quantify mean polyp production along a continuous and non-negative scale, whilst simultaneously modelling the probability of a zero entry (Rigby & Stasinopoulos, 2005). The possibility of a zero entry is necessary because, in *A. aurita*, asexual propagation occurs via multiple pathways, namely lateral budding, and stolon formation (Berrill, 1949). Thus, during propagation, it is possible for a particular pathway to yield no polyps. To obtain a measure of uncertainty for each demographic rate, we also retained the standard deviation of each estimated parameter.

Next, we estimated characteristics associated with the survival and growth of *A. aurita* medusae (Table S1). We modelled medusae survival as a discrete probability, influenced only by temperature and salinity, using a logistic GLMM. We acknowledge that ignoring medusae size likely results in a misrepresentation of survival in very large and very small individuals. Yet, due to their nomadic existence, little is known about the survival of jellyfish medusae (Ceh et al., 2015) and thus, across the literature, data linking medusae size and mortality is limited. We modelled medusae growth and shrinkage using a continuous polynomial relationship between initial medusae size and expected medusae size after a 30-day growth period (necessitated by the available data within the GoJelly databank), with the variance in expected size modelled as a function of initial medusae size, using a gamma distribution (note there was insufficient data to include abiotic factors as fixed effects). We calculated expected medusae size by extrapolating records within the databank linking medusae size and instantaneous growth rate (mm day^-1^). The maximum size medusae could attain during any monthly interval was then defined using a relationship between temperature and maximum observed sizes recorded within the databank; a relationship that we modelled using a gamma distribution in order to ensure a non-negative scale.

We characterised the reproductive output of *A. aurita* individuals using sex ratio, size at maturity, and larval production as a function of size (Table S1). *A. aurita* displays gonochorism and, although variable, the ratio of male to female individuals is often observed to favour females (Eckelbarger & Larson, 1988; Pitt & Kingsford, 2000). In our demographic models we define females to comprise 60% of the population, to correspond with records from *Aurelia* spp. populations in Kiel Fjord, Germany (Javidpour pers. comm. 2021). We then modelled size at maturity as the relationship between mature female medusae size and temperature, set to a gamma distribution. Lastly, we estimated larval production as a function of mature medusae size. For this estimate, data were only available linking the quantity of larvae produced with the wet weight of female medusae (g). Accordingly, we determined the relationship between medusae wet weight and larval output using a gamma GLMM. Importantly, the available data describing larval output represented annual, rather than monthly, contributions, and thus, it was necessary that we scaled these counts to correspond with the monthly timeframe of our modelling framework. To incorporate larval output into our modelling framework, we then relied on the standardised relationship between weight and size (Eq. 4) to convert medusae size (*L*) into a measure of medusae wet weight (*W*).

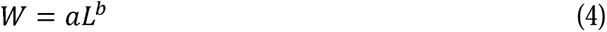

To use this relationship, we extracted the mean and variance estimates of the size and weight coefficients (*a* & *b*) recorded across various *A. aurita* populations and concisely summarised in Hirst & Lucas (1998).

Finally, in some instances, relying solely on the GoJelly databank data proved insufficient for parameterising necessary demographic vital rates. To address this limitation, we enhanced our demographic model by incorporating data from additional sources. For instance, as described earlier, we supplemented our model with information from Hirst & Lucas (1998) describing the relationship between medusae size and wet weight. We used additional data to describe the monthly survival probability of released planula larvae (0.006; Goldstein & Steiner, 2020), and the survival probability of ephyrae undergoing metamorphosis into medusae (0.0095; Ishii et al., 2004). In addition, we also used auxiliary data to describe the physiological limits of medusae survival and reproduction. Irrespective of how temperature and salinity combine to influence the dynamics of *A. aurita* populations, there are certain environmental conditions that remain unsuitable for the establishment of medusae populations, and therefore bloom events (Kennerley et al., 2021). Consequently, corresponding with the environmental and physiological thresholds of *A. aurita* outlined in Conley & Uye (2015), Lucas (2001), and Widmer et al. (2016), we introduced minimum temperature and salinity thresholds into our demographic models, below which medusae survival (≤-1°C & ≤14ppt) and/or reproduction (≤13°C & ≤15ppt) are inhibited.

#### 2.2.2. Implementing forecasting

The second component of our framework, *JellySim*, combines both demographic and abiotic information to implement simulations of the development and spatial movements of the focal jellyfish population. *JellySim* also serves as the user interface for setting up and initiating simulations. The arguments of the *JellySim* function (summarised in Table 1) include user-specified parameters defining the environmental, demographic, and geographic details of the simulation. Firstly, users must specify the geographical extent of their simulations by defining a bounding box with GPS coordinates (*xmx* and *xmn*, for maximum and minimum longitude; *ymx* and *ymn* for maximum and minimum latitude). Within this area, users identify ephyra release points using the *rel_location* argument, which correspond to locations informing the hydrodynamic pathways of drifting individuals (see *2.2.3. Hydrodynamic drifting simulations*), and represent the starting point(s) for subsequent bloom forecasts. Next, users specify the month(s) during which ephyra can occur (*rel_month*), based on suitable abiotic conditions. This argument is useful because ephyra release is mediated by seasonal abiotic patterns and its exact timing can vary interannually (Goldstein & Steiner, 2020; Lucas, 2001). Therefore, the *rel_month* argument allows exploration into how phenological shifts can affect the dynamics of jellyfish blooms.

**Table 1.** Summary of the arguments comprising the *JellySim* function, detailing their roles in configuring simulations of jellyfish population dynamics and environmental interactions.

| Argument Name | Details |
| --- | --- |
| *rel_location* | Specifies the locations within the study area from where jellyfish ephyra are to be initially released. |
| *xmx* | Defines the eastern boundary of the simulations spatial extent (Maximum longitude). |
| *xmn* | Defines the western boundary of the simulations spatial extent (Minimum longitude). |
| *ymx* | Defines the northern boundary of the simulations spatial extent (Maximum latitude). |
| *ymn* | Defines the southern boundary of the simulations spatial extent (Minimum latitude). |
| *rel_month* | Specifies the month(s) during which ephyra release can occur, considering abiotic conditions. |
| *driftData* | Inputs spatial matrices containing abiotic and GPS data describing the movements of, and environmental conditions experienced by, drifting jellyfish. |
| *n_days* | Defines the temporal resolution assumed by the drifting simulations. |
| *pars* | Provides regression coefficients describing the demographic patterns comprising survival, growth/development, and reproduction. |
| *m* | Specifies the dimensions of the demographic projection matrices |
|  | used in the simulation. |
| *zmax* | Sets the number of resampling iterations to account for stochasticity in the forecasts. |
| *parallel* | Determines whether to optimize multicore processing to handle more demanding simulations. |

The *JellySim* function also requires users to provide demographic and environmental parameters for the simulation. The *driftData* argument allows users to input spatial matrices comprising abiotic and GPS records describing the movement and environmental experiences of drifting individuals. Additionally, users must provide *JellySim* with the temporal resolution of these data using the *n_days* argument to define the number of days per month assumed within the drifting simulations (see *2.2.3. Hydrodynamic drifting simulations*). Next, the *pars* and *m* arguments are used to provide regression coefficients describing the demographic patterns of survival, growth/development, and reproduction in their focal population (*pars*), and MPM dimensions (*m*). These two arguments are critical because they enable *JellySim* to align all abiotic and demographic components; thus, they must correspond with the input expectations of the *DMat* function, which is initiated internally by the *JellySim* function. Users can control the extent of variance implemented within the simulations using the argument *zmax* to specify the number of resampling iterations used to account for forecast stochasticity. Users may also select whether to optimise multicore processing for more demanding simulations using the argument *parallel*. With each of these arguments satisfied, *JellySim* initiates the simulations by passing the relevant details to the function *DMat* to determine temporal and spatial patterns in medusae density, which it returns as a series of spatial raster files that can be visualised using the function *SimPlot*.

#### 2.2.3. Hydrodynamic drifting simulations

Although jellyfish can orientate themselves relative to ocean currents to actively influence their spatial movements (Fossette et al., 2015), oceanic currents play a crucial role in mediating spatial patterns in the distribution and aggregation of jellyfish populations (Lehmann & Javidpour, 2010; Schneider, 1987). Therefore, to simulate the spatial and temporal movements of drifting *A. aurita* individuals, within our demonstration, we used particle tracks obtained from simulations of oceanic currents within the Baltic Sea using the ocean modelling system SINMOD (Slagstad et al., 2011; Slagstad & McClimans, 2005; Wassmann et al., 2006).

SINMOD is a three-dimensional model representing the physical (e.g., hydrodynamic, ice dynamics) properties of oceanographic circulation (Slagstad & McClimans, 2005). Here we configurated SINMOD with a 12 km model setup for Nordic and Arctic Seas, with specified boundary conditions covering the Baltic Sea region. Specifying these boundary conditions involved incorporating 12 tidal components imported from the TPXO tide model (https://www.tpxo.net; Egbert & Erofeeva, 2002), and interpolated ERA5 atmospheric data (3 hourly wind forcing, air pressure, daily air temperature, humidity, and cloud cover) from ECMWF (for further details see Hersbach et al., 2020). Riverine input data was also used to specify daily freshwater discharge within the model. For Norwegian rivers, we extracted discharge data from the HBV-model at a 1 km horizontal resolution (Beldring et al., 2003) while all other European riverine input data was sourced from SMHI Hypeweb (https://hypeweb.smhi.se; Arheimer et al., 2020; Lindström et al., 2010). Following configuration, SINMOD uses a Langrangian module to calculate the drift of released particles. To achieve this SINMOD utilises a Runge–Kutta 4th order computational scheme and interpolated gridded 3D velocity fields. Extracting the resultant particle track pathways from SINMOD enabled us to incorporate the spatial movements of drifting *A. aurita* individuals within our predictions of bloom occurrence, alongside sequences of the temperatures and salinities that medusae would be exposed to whilst undertaking these spatial movements. These temperature and salinity sequences were subsequently used by the *DMat* function to determine the abiotic dependant characteristics underpinning the demographic component of our predictive model.

The particle tracks we obtained from SINMOD represent the spatial pathways followed by drifting jellyfish medusae released along coastlines within the Baltic Sea. We defined 1196 sites encircling the Baltic Sea, the Gulf of Finland, the Skagerrak & Kattegat, and the southeast North Sea, to represent potential locations of *A. aurita* polyp populations and subsequently reflect possible release points for drifting medusae (Fig. 3a). We did not define sites within the Gulf of Bothnia, with abiotic conditions here known to inhibit the development of *A. aurita* polyps (Holst & Jarms, 2010). For each location, we applied the hydrodynamic module of SINMOD to replicate the horizontal advection pathways of particles (individual *A. aurita*) released daily throughout 2021 (Fig. 3b). *A. aurita* is primarily observed at depths shallower than the halocline (0–10 metres; Barz & Hirche, 2005; Margonski & Horbowa, 1994). Thus, *A. aurita* particles that were advected or mixed to depths below the halocline (defined as the depth of the first maximum in the vertical salinity gradient, *sensu* Väli et al., 2013) were set to migrate upwards.

**Figure 3.**
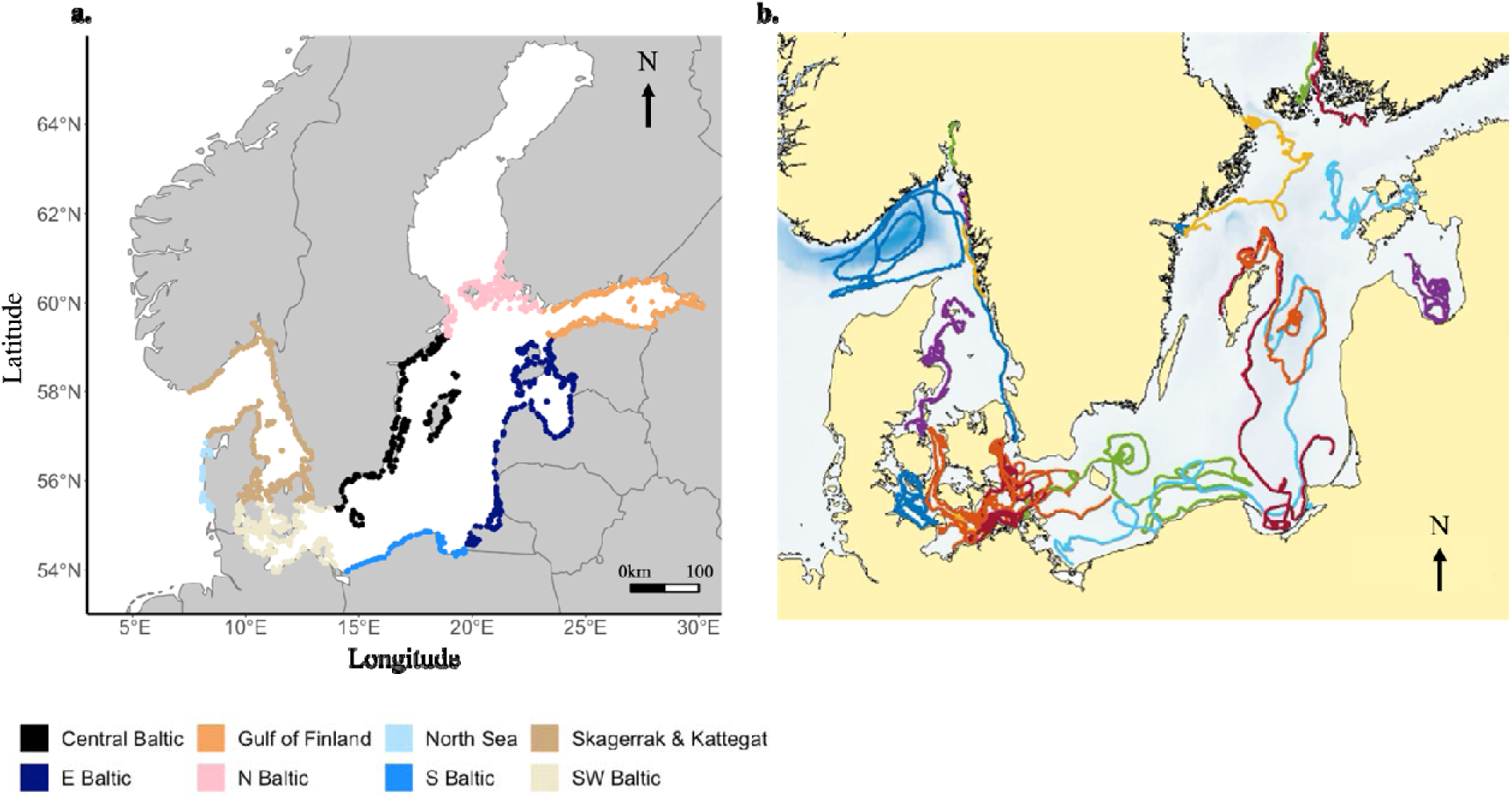
**(a)** 1196 sites defined around the Baltic Sea, the Gulf of Finland, the Skagerrak & Kattegat, and the southeast North Sea, serving as positions of initial particle release within our SINMOD hydrodynamic drifting simulations. These sites were selected to represent potential locations of *Aurelia aurita* polyp populations (*i.e*., the release locations of drifting medusa) within our modelling framework. **(b)** Particle tracks (coloured) extracted from SINMOD, simulating the horizontal movements of particles released daily throughout 2021 from each of our selected release locations. The colour scheme used in panel b does not correspond with the colour key used in panel a, and is only intended to help visualise the diverse range of drifting pathways experienced by particles released across the Baltic Sea.

## 3. Results

Here we illustrate the potential for our modelling approach to predict the spatiotemporal dynamics of jellyfish populations. To demonstrate our framework, we specified the spatial arguments supplied to *JellySim* to focus on forecasting *A. aurita* blooms within the Baltic Sea (Latitude: 53.0–66.0°N; Longitude: 3.0–31.0°E), with ephyra release occurring between February and May (Loveridge et al., 2021; Lucas, 1996). We selected four initial release locations, one from each of the central Baltic, the eastern Baltic, the Gulf of Finland, and the Skagerrak & Kattegat regions. Additionally, we set the resampling count (*zmax*) at 10, the number of days per month (*n_days*) at 30, and demographic projection matrix dimension (*m*) at 200.

The outputs from this forecast consist of maps illustrating spatial patterns in the mean monthly likelihood of encountering *A. aurita* medusae (Fig. 4), and the confidence of these forecasted likelihoods (Fig. 5). Across this forecast, medusae density is presented as a proportional measure of the number of medusae produced per polyp (med polyp^-1^) rather than medusae density per unit volume of seawater. This approach was necessary because the initial population structure is defined within our framework using polyp density per unit area, which is then used to estimate a count of the number of individual ephyra released. Data on the spatial extent of polyp populations within the Baltic Sea is almost non-existent and there is little evidence for how the number of ephyra released at a particular location relates to medusae development per unit volume of seawater. Therefore, it was not possible to rescale our initial population structure into medusae density per unit volume, nor estimate absolute medusae counts. By using proportional medusae densities instead, we ensure that the relative temporal and spatial patterns in medusae dynamics reported by our model are still valuable for predicting the location and timing of jellyfish aggregations.

**Figure 4.**
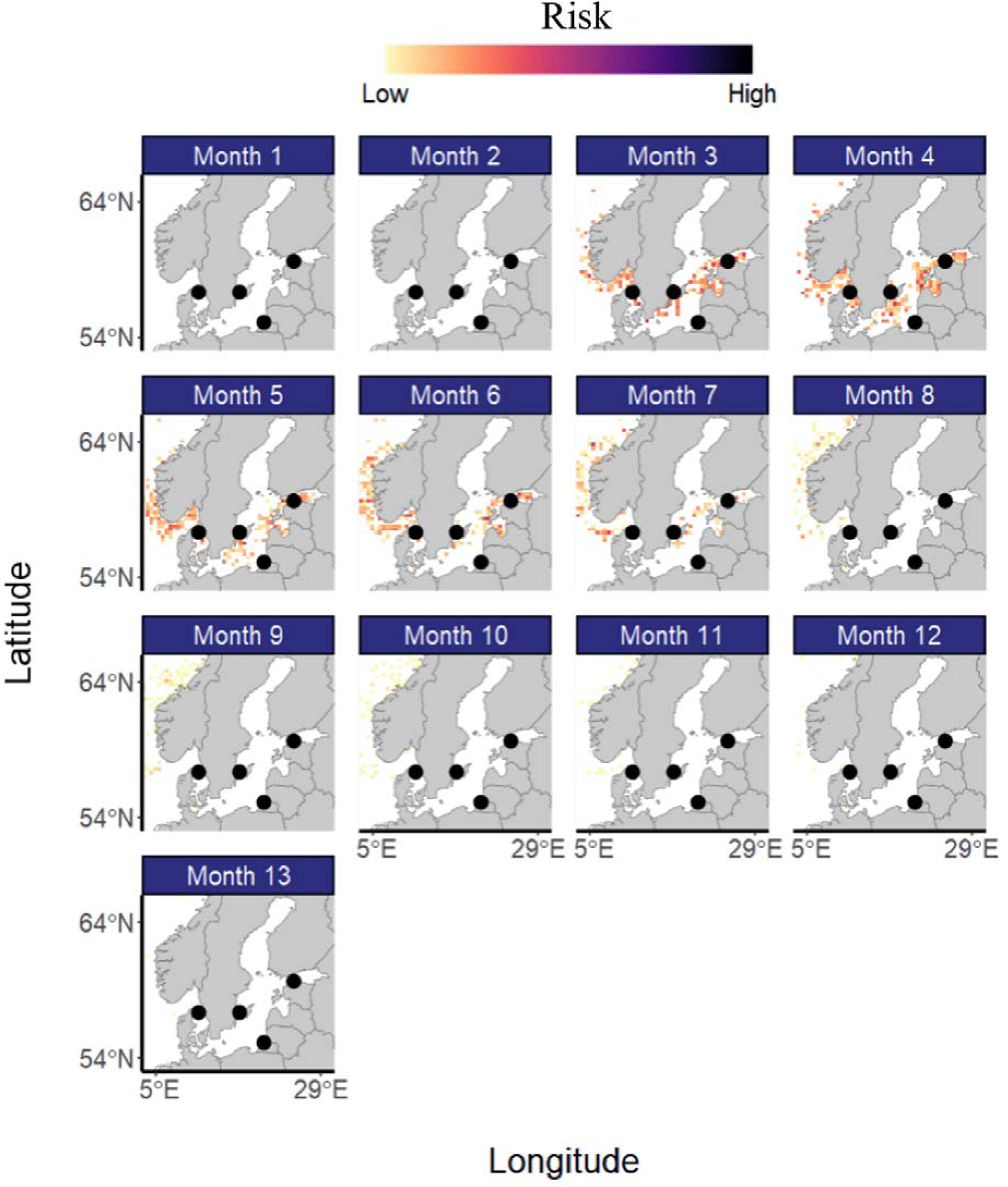
Spatio-temporal patterns in predicted *Aurelia aurita* medusae densities within the Baltic Sea following their release from polyp beds located in the Skagerrak & Kattegat region (A), the central Baltic (B), the Gulf of Finland (C), and the eastern Baltic (D). Patterns in Medusae density is displayed using a colour scale representing the likelihood (‘Risk’) of encountering medusae in a given region. These forecasts were generated using daily records describing the spatial movements of drifting particles released within the Baltic Sea, averaged over a 30-day month period. Subsequently, it is possible for individuals released during the first 5 days of the year to experience 13 ‘months’ over a 365-day period.

**Figure 5.**
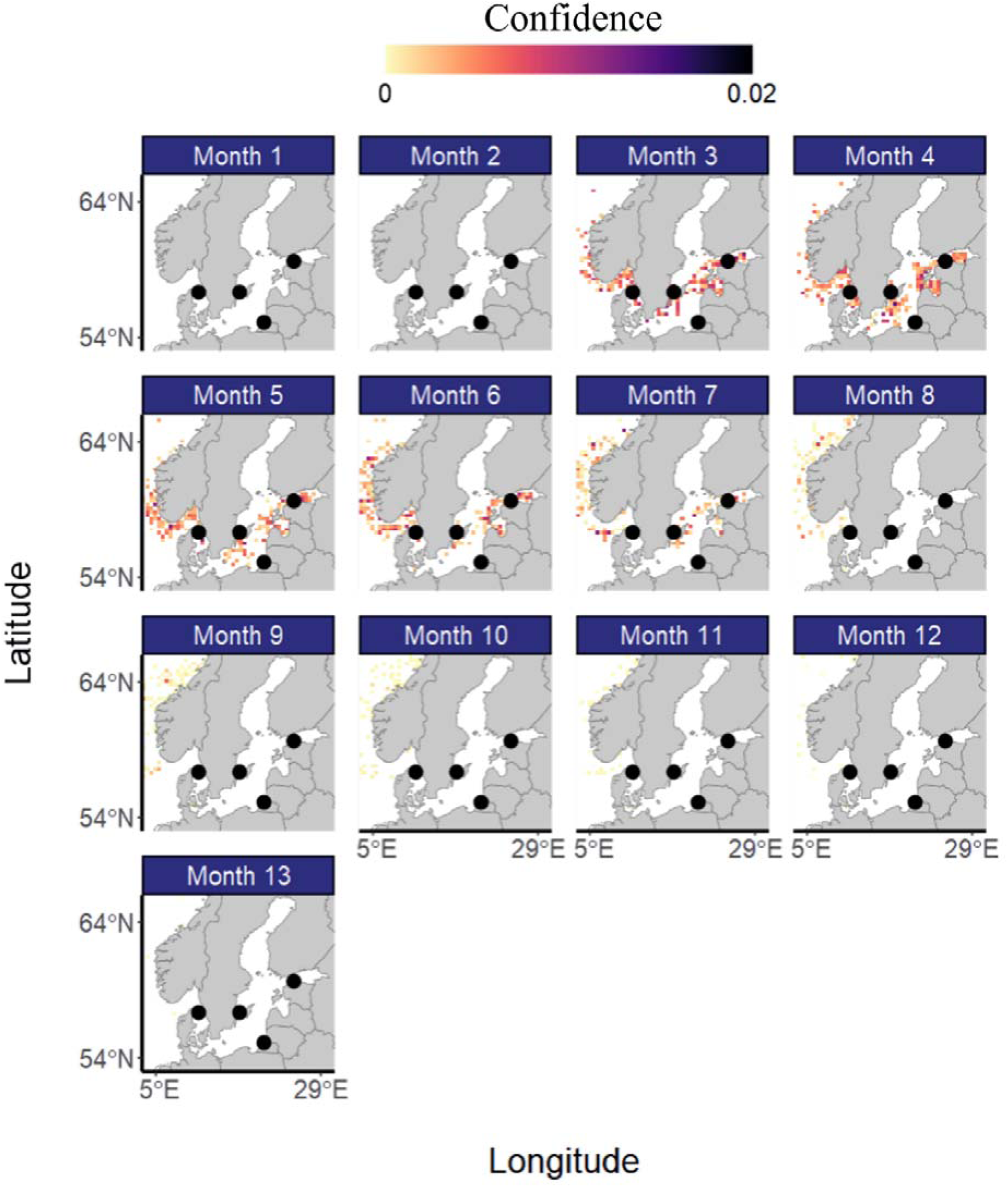
Spatio-temporal patterns in the confidence associated with predicted *Aurelia aurita* medusae densities within the Baltic Sea following their release from polyp beds located in the Skagerrak & Kattegat region (A), the central Baltic (B), the Gulf of Finland (C), and the eastern Baltic (D). Confidence estimated as the standard deviation of medusae densities calculated across 10 iterations of the initial scenario accounting for variation in exact release day during any given month and demographic stochasticity. These forecasts were generated using daily records describing the spatial movements of drifting particles released within the Baltic Sea, averaged over a 30-day month period. Subsequently, it is possible for individuals released during the first 5 days of the year to experience 13 ‘months’ over a 365-day period.

In our illustrative scenario, *A. aurita* medusae are expected to first appear in early spring, with their densities peaking a month later, particularly in the Gulf of Finland and around the coastlines of Estonia and Latvia (Fig. 4). Throughout the summer, *A. aurita* densities are then expected to decline within the Baltic Sea, whereas in the North Sea abundances remain more consistent through the summer with the population then drifting northward along the coast of Norway as it diminishes through the autumn and winter (Fig. 4). Contrastingly, despite possessing an initial release location, coastal regions in the southern and south-eastern Baltic Sea will likely experience very few *A. aurita* aggregations (Fig. 4). While the accuracy of predicted population densities tends to decrease with higher predicted values, the standard deviation of the majority of predicted densities remains constrained below 0.01 (Fig. 5).

## 4. Discussion

As ongoing climatic change and a growing human population place increasing pressure on our natural environment and alter the dynamics of human-wildlife interactions, we are challenged to improve our capacity for predicting the movements and abiotic responses of biological assemblages (Evans, 2012). Here, we have presented a novel framework for predicting spatio-temporal patterns in the movements and aggregation of jellyfish populations to elevate our capacity to forecast the occurrence of jellyfish bloom formation. Our approach represents a form of agent-based modelling (ABM), which considers the ‘decision-making’ process of individuals in refining where, and when, they move (DeAngelis & Mooij, 2005). Occurring both consciously and unconsciously (invariably the latter in the case of jellyfish) in response to environmental conditions, individual decision-making underpins population performance and viability (Beekman & Jordan, 2017). Yet, despite this value, the challenges associated with empirically modelling the mechanisms underlying individual ‘choice’, means that ABM tools are often overlooked during the development of conservation and management strategies (DeAngelis & Diaz, 2019). Accordingly, our approach here advocates for enhancing the implementation of ABM tools to improve our capacity to mitigate the socioeconomic impacts of jellyfish blooms, and represents the beginning of efforts to quantify the cryptic spatial dynamics of global jellyfish populations.

Empirical records as to how the transition of jellyfish individuals through their complex and cryptic lifecycles influences the spatial dynamics of their populations do not currently exist at the resolution necessary for quantitatively predicting the exact timing and location of jellyfish bloom occurrences (Fernández-Alías et al., 2024). This lack of capacity is largely because identifying jellyfish polyp beds is imperative for understanding the subsequent movements of medusae aggregations, and yet these seed locations remain frustratingly mysterious (Gibbons & Richardson, 2013). Indeed, resolving this knowledge gap was a key motivation behind our development of this framework. The statement ‘*all models are wrong, but some are useful*’ (Box, 1979) captures the idea that we cannot hope to reflect the inherent complexity of natural systems using analytical models. However, we have designed our framework to help enhance our ability for identifying potential seed locations using the spatial and temporal details of known bloom events; thereby highlighting regions of high priority for conducting future polyp-bed surveys. Citizen science initiatives (e.g., Jelly Spotter & Medjelly) are being increasingly used to record the location and abundance of observed jellyfish individuals and aggregations (Johansen et al., 2021; Marambio et al., 2016). Coupled with the increased use of satellite and drone technology in jellyfish monitoring (Mcilwaine & Rivas Casado, 2021), these approaches offer real-time information on the appearance of jellyfish blooms. With our framework predicting the movements of drifting jellyfish individuals following their release from user defined release sites, validating the extent to which these forecasted patterns correspond with real-time observations offers an approach for revealing potential polyp bed locations. Identifying the locations of hitherto unknown polyp beds will be a game-changer in our ability to predict jellyfish blooms and manage their economic impacts. Consequently, our framework, holds potential for becoming a valuable decision-support tool for guiding the future management of jellyfish populations.

It is important to acknowledge here that, due to the challenges associated with tracking the location of polyp beds and the settlement of planula larvae produced by drifting medusae, it is not possible for our predictive framework to consider the future contribution of second-generation individuals towards bloom formation. Accordingly, our framework currently only forecasts over a single annual interval and cannot yet be used to explore the multiyear dynamics of bloom occurrence. Again, this framework and the steps involved in its parameterisation are intended to explicitly highlight key knowledge gaps hindering the prediction of the spatial dynamics of jellyfish aggregations. However, we have intentionally designed this framework to be adaptable as and when new information becomes available. Accordingly, rates of larval production are still parameterised within the demographic component of our model to facilitate the future inclusion of their settlement dynamics. Nevertheless, given the critical importance of this life stage, future research should focus on identifying polyp bed locations, which could significantly improve the predictive capabilities of our model.

The predictive tool we present here is also intended to increase the wider public understanding of jellyfish ecology and the spatial movements of their populations. Increasing public engagement in the occurrence of jellyfish blooms is necessary for the successful implementation of strategies mitigating their economic impacts, and their conservation, and will help to improve the quantity and quality of observer data available for refining future modelling efforts (Doyle et al., 2014). Thus, the methodology we described above, coupling demographic and hydrodynamics drifting simulations, has also been incorporated into an interactive jellyfish bloom risk map (https://gojellyeu.shinyapps.io/gojellyapp/). We created this application, by collating the analytical functions, *JellySim*, *DMat*, and *SimPlot*, into an accessible user interface using the *R* package *Shiny* (Chang et al., 2021). Consistent with our illustrative example, this interactive map currently focuses on forecasting and visualising the spatial movements of *A. aurita* populations within the Baltic Sea. However, like our overall framework this application is sufficiently flexible so that it can be scaled up, both geographically and taxonomically, to accommodate other regions impacted by problematic jellyfish blooms (e.g., the Mediterranean Sea [Brotz & Pauly, 2012] and South-East Asia [Syazwan et al., 2020]), and other jellyfish species. The demographic inputs required by our framework comprise a series of regression coefficients outlining the vital rates of the focal population(s). Provided the demographic modelling function, *DMat*, is adapted to accommodate these variables, then the rest of the framework will adjust accordingly. Moreover, provided the model is supplied with the appropriate GPS coordinates corresponding with any simulated movements, the hydrodynamic simulations provided to the framework describing the spatial movements of drifting individuals, can be changed to reflect ocean current pathways in various regions across the globe.

### 4.1. Conclusions

Jellyfish possess a largely negative public image underpinned by reports of fatal stings, fisheries obstruction, and the blocking of electrical supplies and cooling systems (Doyle et al., 2014; Gershwin et al., 2013; Graham et al., 2014). Mitigating the negative human-jellyfish interactions, and managing the economic impacts, arising from these encounters requires a detailed understanding of their spatial movements and spatio-temporal patterns in the occurrence of jellyfish blooms. Concurrently, apparent increases in the reoccurrence of jellyfish blooms (although yet to be formally substantiated), further emphasises the urgent need for an ability to forecast the location(s) and timing(s) of jellyfish blooms. Regrettably, however, we currently lack sufficient knowledge of the spatial dynamics of jellyfish populations in order to achieve this level of understanding. Here, we combine state-of-the-art drifting simulations with a comprehensive synthesis of existing knowledge regarding the ecological dynamics of a cosmopolitan jellyfish species, to develop a novel framework for forecasting the dynamics of jellyfish bloom occurrence. A framework that, combined with iterative validation through field observations, will help us to identify the locations of jellyfish polyp beds. By embracing the concept of using models and refining them over time to both identify and resolve knowledge gaps (Dietze & Lynch, 2019; Houlahan et al., 2017), such a framework will be beneficial for enhancing our understanding of the cryptic processes underpinning the formation of jellyfish blooms.

## Author Contributions

JJ, ORJ, NA, SM, and JD devised the initial project idea. JJ, NA, and SM sourced project funding. JC led data curation, developed the modelling framework, and created the webapp, with guidance from ORJ. JC, ORJ, and JL implemented all analyses. IE carried out all hydrodynamic drifting simulations. JC led manuscript write up with all authors contributing to subsequent drafts.

## Acknowledgements

This project was funded by the European Union‘s Horizon 2020 research and innovation program (Grant agreement no. 774499) as part of GoJelly (work package 2: ‘Driving mechanisms and predictions of jellyfish blooms’).

## Conflict of interest statement

The authors declare that there are no conflicts of interest associated with this work.

## Data availability

All modelling code described throughout this manuscript is publicly available on GitHub (https://github.com/CantJ/Bloom-forecasting-tools), and will be archived on Zenodo once this manuscript is accepted for publication. Meanwhile, further details on the SINMOD ocean model used in this study can be found on the SINTEF website (https://www.sintef.no/en/ocean/initiatives/sinmod/).

## Notes

### Competing Interest Statement

The authors have declared no competing interest.

