## Supplementary Material for "Coupling hydrodynamic drifting simulations and seasonal demographics to forecast the occurrence of jellyfish blooms"

**Quantifying the demographic characteristics underpinning patterns in individual survival, growth, and reproduction, across the life cycle of *Aurelia aurita*.**

**Table S1.** Using the GoJelly databank (Volpe et al., 2023) we calculated a series of demographic characteristics underlying patterns of survival, growth, and reproduction across the planula, scyphistoma, ephyra, and medusa phases of the *Aurelia aurita* life cycle. Here each demographic characteristic is displayed alongside colour scales depicting whether the characteristics (1) reflect a discrete or continuous transition (Light grey = discrete; Dark grey = continuous), (2) were estimated as environmentally explicit (Navy = influenced by both temperature and salinity; Light blue = only temperature; No colour = No abiotic influence), and (3) were estimate using only data extracted from the Biobank (No colour = True; orange = additional sources used).

| **Life-stage** | **Demographic characteristic** |
| --- | --- |
| Planula | % Settlement |
|  | % Survival |
| Scyphistoma | % Survival |
|  | % Strobilation |
|  | No. ephyra produced polyp^-1^ |
|  | % Asexual propagation |
|  | No. polyps produced polyp^-1^ |
|  | Polyp density (Polyp cm^-2^) |
| Ephyra | % Survival |
|  | % Metamorphosis |
| Medusae | % Survival |
|  | Size transition |
|  | Maximum size |
|  | Size at maturity |
|  | Sex ratio |
|  | No. Larvae produced ~ Wet weight |
|  | Size/Weight relationship |
